# Essential bromodomain *Tc*BDF2 as a drug target against Chagas disease

**DOI:** 10.1101/2022.01.25.477728

**Authors:** Alejandro Pezza, Luis E Tavernelli, Victoria L Alonso, Virginia Perdomo, Raquel Gabarro, Rab Prinjha, Elvio Rodríguez Araya, Inmaculada Rioja, Roberto Docampo, Felix Calderón, Julio Martin, Esteban Serra

## Abstract

*Trypanosoma cruzi* is a unicellular parasite that causes Chagas disease, which is endemic in the American continent but also worldwide distributed by migratory movements. A striking feature of trypanosomatids is the polycistronic transcription associated with post-transcriptional mechanisms that regulate the levels of translatable mRNA. In this context, epigenetic regulatory mechanisms have been revealed of great importance, since they are the only ones that would control the access of RNA polymerases to chromatin. Bromodomains are epigenetic protein readers that recognize and specifically bind to acetylated lysine residues, mostly at histone proteins. There are seven coding sequences for BD-containing proteins in trypanosomatids, named *Tc*BDF1 to *Tc*BDF7, and a putative new protein-containing a bromodomain that was recently described. Using the Tet regulated overexpression plasmid p*Tc*INDEX-GW and CRISPR/Cas9 genome editing we were able to demonstrate the essentiality of *Tc*BDF2 in *T cruzi*. This bromodomain is located in the nucleus, through a bipartite nuclear localization signal. *Tc*BDF2 was shown to be important for host cell invasion, amastigote replication, and differentiation from amastigotes to trypomastigotes. Overexpression of *Tc*BDF2 diminished epimastigote replication. Also, some processes involved in pathogenesis were altered in these parasites, such as infection of mammalian cells, replication of amastigotes, and the number of trypomastigotes released from host cells. In *in vitro* studies, *Tc*BDF2 was also able to bind inhibitors showing a specificity profile different from that of the previously characterized *Tc*BDF3. These results, point to *Tc*BDF2 as a druggable target against *T. cruzi*.

---

The World Health Organization (WHO) estimates the prevalence of Chagas’ disease (or American Trypanosomiasis) in about 7 million cases, of which between 20 to 30% develop severe clinical manifestations ^1,2^. Originally confined to Latin America, migration is spreading the disease to non-endemic countries, where it can be transmitted by blood transfusion, organ transplantation, and from infected women to their offspring during pregnancy. There are two active compounds (nifurtimox and benznidazole) that are used in the treatment of the acute phase of the disease. Both have severe side effects and their efficacy in the chronic phase is controversial. This situation has made evident the need to find new trypanocidal compounds that can be used as therapeutic alternatives for all stages of the disease ^3^.

In Trypanosomatids, genes are organized and transcribed in a quite unusual polycistronic fashion. The primary long transcripts are processed to mature translatable mRNAs by adding a 5’capped common short RNA sequence by *trans*-splicing and a 3 poly-A tail ^4^. The final gene expression levels are regulated by several post-transcriptional mechanisms, which are not yet completely understood ^5,6^. In parallel, transcription appears to be controlled by several proteins, histones and non-histones, which take part in protein complexes that regulate chromatin accessibility and are now starting to be identified ^7–9^. Chromatin structure can be altered by post-translational modifications in the N-terminal tails of histones, by the substitution of one or more histones by their variants, or by combinations of these events ^10^. These conformational changes regulate not only the ability of chromatin to be transcribed but also other DNA-related processes such as recombination, replication, repair, and chromosomal segregation during metaphase ^11^.

Regarding post-translational modifications of histones, acetylated H3 and H4 have been mapped to the beginning of the polycistronic units, considered to be the RNA Polymerase II (RNAPII) transcription initiation sites in *T. cruzi, Trypanosoma brucei,* and *Leishmania* spp, suggesting a conserved localization for canonical histones among the TriTyps ^12–14^. Histone variants have been also mapped in *T. brucei* and *Leishmania major* ^15–18^. H2A.Z and H2B.V, usually associated with less stable nucleosomes, were found to be enriched at the putative RNA Pol II transcription start regions (TSRs), while histone variants H3V and H4V (that seems to be specific of *T. brucei*) were found enriched at the end of the polycistronic transcription units, suggesting a role in transcription termination^15–19^. Acetylated H2A.Z at the TSRs was also associated with the transcription level of the whole gene tandems ^20^. More recently, H2B.V was found at the beginning of gene clusters in divergent strand switch regions in *T. cruzi,* as well as in some tDNA loci ^21^. Interestingly, *Tc*H2B.V was also found between core and disruptive genome compartments. In these zones, multigenic families of surface proteins like *trans*-sialidases, MASP and mucins, are concentrated ^22^.

Bromodomains (BD) form a family of “modifier reader” proteins that recognize the acetylation marks on histones working as a scaffold for the assembly of macromolecular complexes that interact with chromatin ^23,24^. New and more complex functions for this protein family have been proposed over the last years ^25^. Some proteins containing BDs are now considered druggable and BD inhibitors are currently assayed in a myriad of pathologies such as cancer or inflammatory diseases among others ^25–28^. Currently, 44 clinical trials concerning bromodomain inhibitors are listed at the ClinicalTrials.gov database from NCBI (https://clinicaltrials.gov/) ^29^. Parasite bromodomains are also proposed as drug targets, and a small number of inhibitors have been previously tested in *T. cruzi, T. brucei, Plasmodium falciparum,* and *Toxoplasma gondii* ^16,30–36^.

Our laboratory has been interested in the characterization of bromodomain factors from *T. cruzi* (*Tc*BDFs). According to the TriTrypDB (http://tritrypdb.org/tritrypdb/) ^37^, there are seven coding sequences for BD-containing proteins in trypanosomatids, named *Tc*BDF1 to *Tc*BDF7. An eighth putative bromodomain factor recently identified in *Leishmania* spp has an orthologue in *T. cruzi ^9^.* Even though a few *Tc*BDFs have recognizable additional domains, the architecture of these proteins seems to be simpler than those of BD-containing proteins from fungi and animals. For example, none of them contain a histone acetyltransferase (HAT) domains, as occurs in the GCN5 acetyltransferase family, which is conserved in almost all eukaryotic cells, including many protists^38^.

The BDs from TriTryp BDFs share limited identity among each other as with any other eukaryotic BDs ^39^. The inner core is different in each domain and only a limited number of the hyper conserved amino acids are present, also the hydrophobic pocket is mostly constituted by non-identical amino acids with conserved hydrophobicity. This huge divergence of BDs from trypanosomatids is not surprising considering the highly atypical nuclear biology of these early branched protists. Despite this, the solved crystallographic structure of *Tc*BD2 and *Tc*BD5.1 (from *Tc*BDF2 and *Tc*BDF5 respectively) showed the typical four alpha helices, each separated by two loops that form the hydrophobic acetyl lysine-binding pocket (Structural Genomics Consortium; PDB ID: 6NP7 and 6NEY). Moreover, new reliable structures were generated by AlphaFold2, showing the typical fold of the domain for all the *T. cruzi* bromodomains (https://alphafold.ebi.ac.uk)^40^. Surprisingly, *Tc*BDF7 one of the most conserved proteins of the family, shows a typical BD folding but the hydrophobic pocket seems to be buried in the predicted structure.

We have previously characterized *Tc*BDF1, *Tc*BDF2, and *Tc*BDF3 from *T. cruzi.* We found that *Tc*BDF1 and *Tc*BDF3 are mainly localized to the cytosol of *T. cruzi* cells. *Tc*BDF1 was found in the glycosome, it is developmentally regulated throughout the *T. cruzi* life cycle, and its importance for parasite fitness has been well established ^41^. *Tc*BDF3, which was found to be associated with acetylated α-tubulin at the subpellicular corset and flagella, was found necessary for the differentiation of both epimastigotes and intracellular amastigotes to trypomastigotes^42^. In addition, complete knockouts for *Tc*bdf1 and *Tc*bdf3 could not be obtained by CRISPR/Cas9 genome editing (unpublished results from our laboratory). We also described two compounds that bind to *Tc*BD3 *in vitro* and also have trypanocidal activity ^30–32^. *Tc*BDF2 was the first BD to be described in trypanosomatids and it is expressed throughout the whole life cycle of the parasite, and located in discrete regions of the nucleus ^43^. In the same report we showed that it can interact with histone H4 through acetylated K10 and K14 and detected interaction with H2 by far-western blot analysis ^43^. Recently, *T*cBDF2 was found to be associated with H2B.V in pull-down assays and it was also shown to interact *in vitro* with H2B.V, H2B, and H4 ^44^.

In contrast to *T. cruzi* BDFs, all BD-containing proteins from *T. brucei* were described to be nuclear, and it has been proposed that many of them are essential for the maintenance of the bloodstream form of this parasite ^45,46^. However, a recent study showed that *Tb*BDF4 has a bipartite nuclear-cytosolic localization ^8^. Even though a genome-wide RNA interference viability screen showed that *Tb*BDF2 was not essential, it was found essential in another assay ^46^. Very recently, Jones *et al.* showed that five BDFs, among them BDF2, are essential for *Leishmania mexicana* promastigotes ^9^.

There are also some discrepancies about the sites of interaction of *Tb*BDF2 within the genome. Early studies did not find *Tb*BDF2 enriched at RNAPII TSRs ^13,46^. However, *Tb*BDF2 was found later to be bound to the hyperacetylated form of the variant histone H2A.Z at the TSRs and placed also at the nucleolus ^16^. Later, in a systematic study of *T. brucei* potential chromatin factors, *Tb*BDF2 was found broadly enriched at TSRs, following the direction of RNAPII transcription for 5-10 kb together with H2A.Z and H2B.V ^8^. In this report *Tb*BDF2 was shown to immunoprecipitate with many proteins with recognizable HDAC, helicase, HIT, and CW-type zinc fingers and histone binding domains, among others. Based on a global interpretation of all their immunoprecipitation results, Staneva *et al.* proposed that *Tb*BDF2 could be included in at least two complexes: (1) a chromatin remodeling complex related to SWR1, which is responsible for the incorporation of H2A.Z into chromatin in other cells, and (2) a more intriguing complex assembled over the telomerase-repeat binding protein (TRF) and then, presumably located at telomers ^8^.

In the present report, we demonstrate that *Tc*BDF2 is essential for the development of both epimastigotes and intracellular amastigotes and validate it as a target for the search of new trypanocidal drugs using commercial bromodomain inhibitors.

## Results and discussion

### *Tc*BDF2 gene (*TcBDF2*) is essential in epimastigotes

In order to investigate how *Tc*BDF2 is involved in different aspects of *T. cruzi* physiology, we tried to generate a *Tc*BDF2 knockout mutant line and evaluate the resulting phenotypes. We followed the CRISPR/Cas9 genome editing strategy reported by Lander *et al* ^47^, which is outlined in Figure 1A. Briefly, the sgRNA targeting *TcBDF2* sequence was cloned into the pTREXCas9 vector and co-transfected into the parasite with the donor DNA. The sgRNA guides the Cas9 complex to the specific site in the DNA and produces a double-strand break in the DNA of the selected gene. Then, this rupture induces the DNA to be repaired by double homologous recombination in the presence of a donor DNA that carries a selectable marker (blasticidin in this case). Once the resistant parasites were selected we isolated the genomic DNA and corroborated donor DNA insertion by PCR (Figure 1B, inset). Despite three attempts to knockout *TcBDF2*, we were only able to obtain parasites with only one mutated copy of the gene, suggesting that *TcBDF2* is an essential gene for epimastigotes. As is shown in Figure 1B we evaluated the proliferation of epimastigotes of the heterozygous line and determined that there were no significant differences in the growth rate when compared to the control line, indicating that one copy of the gene is enough for epimastigotes’ normal growth.

**Figure 1.**
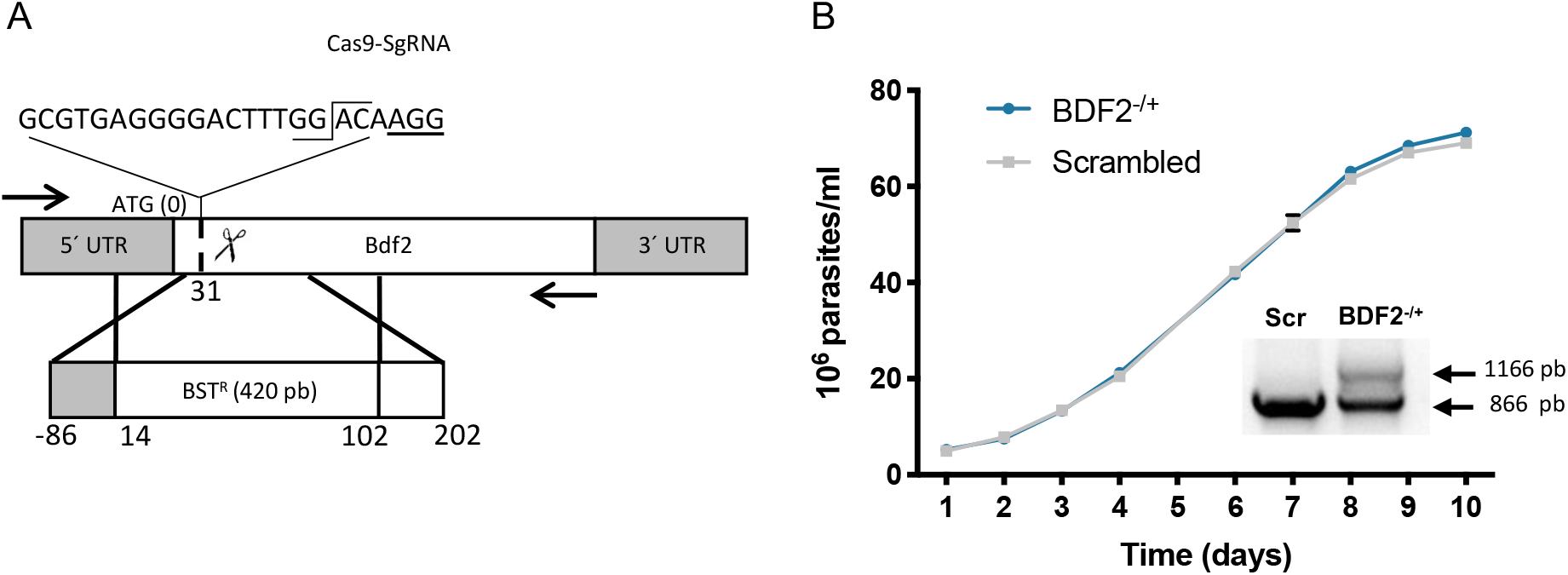
(A) Schematic representation of *TcBDF2* mutation by CRISPR/Cas9 genome editing. The whole strategy was designed to replace the *TcBDF2* coding sequence with a blasticidin resistance gene (BST^R^). Taking the adenine of the initial codon as zero, numbers indicate the start and end of each homology region. The protospacer sequence and cutting site position are also indicated. The arrows represent the oligonucleotides employed to determine the presence of the mutation. (B) Growth curve of epimastigotes transfected with scramble sgRNA (black squares, Scr) or BDF2^-/+^ (blue circles) cells, counted daily for 10 days. The KO allele generates an 1166 pb PCR product while the WT PCR product size is 866 pb with the oligonucleotides indicated in A (inset (B)).

### *Tc*BDF2 bears a functional canonical NLS within its C-terminal domain

As mentioned before, several attempts to establish a stable KO line by CRISPR/Cas9 were carried out and despite our effort, we were not able to knockout *TcBDF2* completely. Therefore, we decided to pursue a different methodological approach to deepen the knowledge of *Tc*BDF2 in the parasite life cycle. We made use of the vector p*Tc*INDEXGW, a GATEWAY® Cloning Technology-adapted version of p*Tc*INDEX^48,49^ built in our laboratory. This plasmid allowed us to overexpress our protein of interest under the control of a tetracycline-regulated promoter in all the stages of the *T. cruzi* life cycle ^48,49^.

First, we constructed different truncated versions of *Tc*BDF2 to determine the localization of its predicted nuclear localization signal (NLS). *In silico* study of *Tc*BDF2 amino acid sequence by NLstradamus ^50^ predicted a complete nuclear localization site (NLS) between the amino acids R162 and N177, which is present in BDF2 from all *T. cruzi* strains, but absent in *Leishmania* spp and *T. brucei.* To test their functionality, we cloned the full coding sequence of *Tc*BDF2 (*Tc*BDF2HA) as well as truncated mutants with (*Tc*BDF2Δ177) or without (*Tc*BDF2Δ162) the largest predicted NLS region, into the plasmid p*Tc*INDEX-GW. These constructs were tagged with an epitope of hemagglutinin (HA) at the N-terminus and used to transfect Dm28c/pLew13 epimastigotes (Fig. 2A).

**Figure 2.**
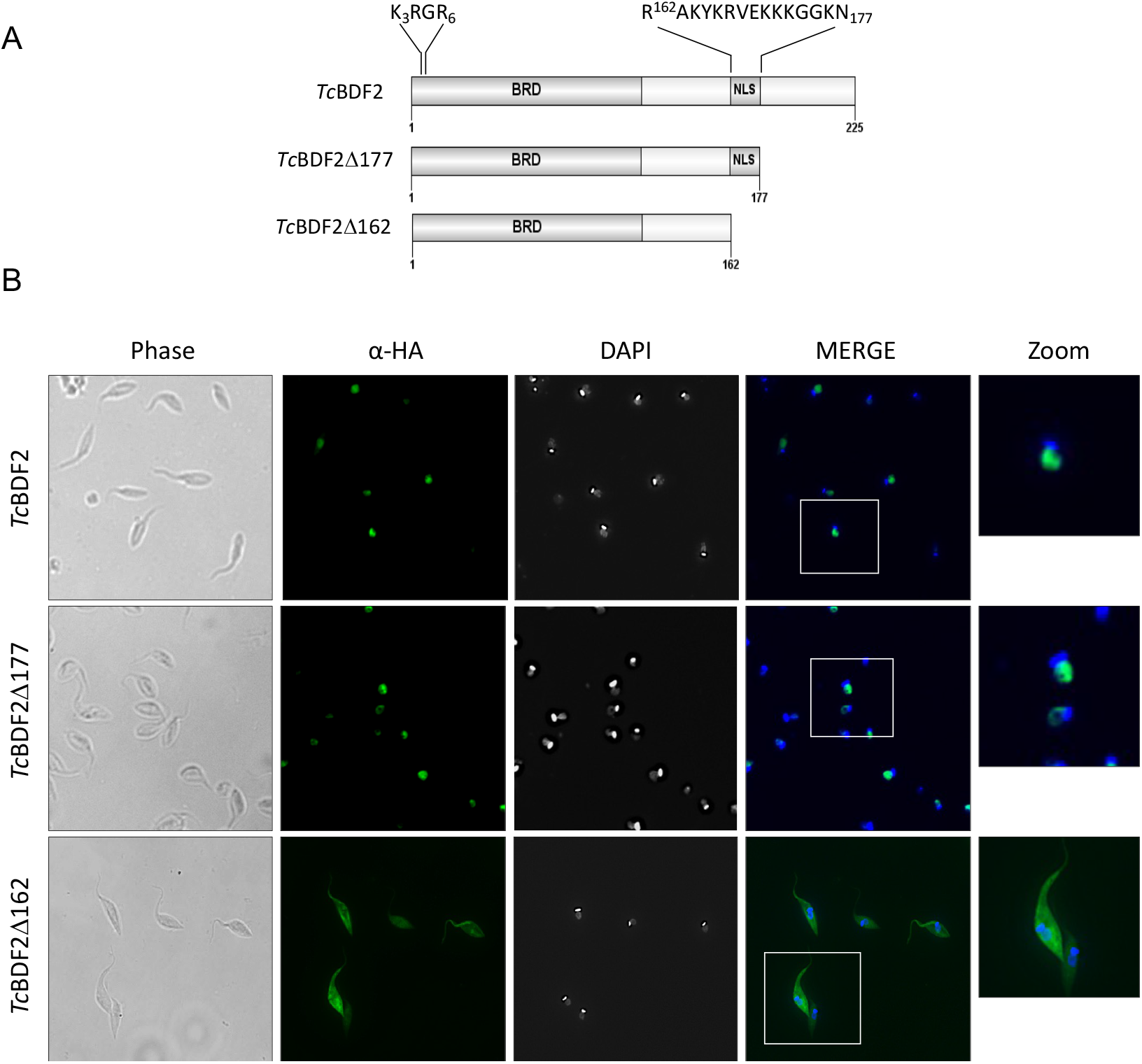
A Schematic representation of the truncated versions of *Tc*BDF2 B. Localization of *Tc*BDF2HA (wild type) and C-terminal truncated proteins (HA-tagged) analyzed by immunofluorescence with anti HA antibodies (α-HA). Images were obtained using monoclonal rat anti-HA antibodies developed by anti-rat IgG antibody conjugated to Alexa 488 (green). The nucleus (N) and kinetoplast (K) were labeled with DAPI. The scheme on the left shows the different mutated versions of *TcBDF2* which expression pattern in epimastigotes is shown on the right. The bromodomain portion of the protein is indicated as BRD and the predicted Nuclear Localization Signal is indicated as NLS. Scale bar = 5 μm.

As expected, *Tc*BDF2HA was directed to the nucleus (Fig. 2B), giving a strong nuclear signal in the epimastigote stage when immunolocalized with anti-HA antibodies, that was conserved when the last 48 amino acids where deleted in the deletion mutant *Tc*BDF2Δ177HA. We have previously reported that *Tc*BDF2 is localized in the nucleus under physiological conditions ^43^. On the contrary, when we overexpressed the mutated version *Tc*BDF2Δ162, a clear change in the localization of the signal from the nucleus to the cytosol was observed. These results clearly show that the NLS predicted is responsible for targeting and retaining *Tc*BDF2 in the nucleus.

It has been proposed that the functional monopartite NLS in *T. brucei* KRxR, a sub-motif of a classical eukaryotic NLS, is the characteristic NLS for trypanosomatids ^51^. When we inspected the NLS predicted region in detail we observed that *Tc*BDF2 shows also a monopartite NLS at the N terminus between the amino acids R3 and K6. This putative signal (KRGR) is well conserved in all *T. cruzi* strains and *Leishmania* spp (KRPR) but absent in the *T. brucei* homolog. However, it is clearly not able to target *Tc*BDF2Δ162 to the nucleus, suggesting that is not functional in *T. cruzi,* as has been proposed by Goos *et al.* for trypanosomatids based in the *T. brucei* nuclear proteome ^51^. Even though further studies will be necessary to determine which specific amino acid/s is/are responsible within the NLS for the localization of *Tc*BDF2 to the nucleus, it is clear that the canonical NLS, present in *T. cruzi* but absent in other trypanosomatids, is one of the many differences that the bromodomain-containing protein family show among trypanosomatids.

### Overexpression of both wild type and mutant *Tc*BDF2 produces a detrimental effect on epimastigote growth

Next, we generated *Tc*BDF2 constructs with two single point mutations that change essential amino acids located at the hydrophobic pocket of the BD. Previously, we demonstrated that the change of two conserved hydrophobic amino acids of *Tc*BDF3 to alanines abolished the capability of the protein to recognize its ligand, but at the same time, the overall protein-folding remains unaltered. Following the same strategy, we identified and selected Y85 and W92 to be substituted by alanine residues (hereinafter *Tc*BDF2HAdm).

*Tc*BDF2dm and *Tc*BDF2 wild type constructs were cloned in the p*Tc*INDEX-GW plasmid with HA tags and were transfected to epimastigotes. This overexpression approach of a mutant BD has been successfully used by our group to generate a dominant-negative phenotype that was unable to bind to its acetylated target ^30,41^.

We corroborated the correct expression of the inducible system by western blot analyses of both *Tc*BDF2HAwt and *Tc*BDF2HAdm. As can be seen in Figures 3 A and B (right panel), we detected only one band of the expected molecular weight when probed with an a-HA antibody. Even though correct folding of *Tc*BDF2HAdm was not verified when expressed in *T. cruzi,* both the wild type and mutant bromodomain were demonstrated to be folded after expression and purification from *E. coli* (see below). *Tc*BDF2HAdm, as well as *Tc*BDF2HA, were also detected in the nucleus when IFA was performed using an a-HA antibody (Supplementary Figure 1).

**Figure 3:**
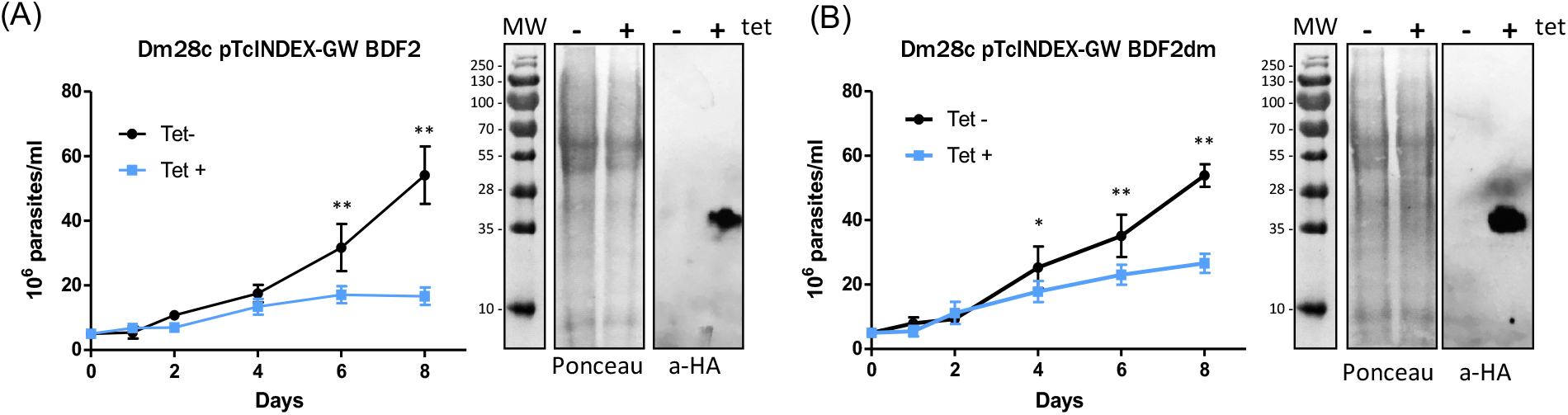
*Tc*BDF2HA and *Tc*BDF2HAdm overexpression decrease cell proliferation. Growth curve of epimastigotes transfected with p*Tc*INDEX-GW-*Tc*BDF2HA (A) or p*Tc*INDEX-GW-*Tc*BDF2HAdm (B) in the absence (blue circles, Tet-) or presence (black squares, Tet+) of 0.5 μg/ml tetracycline, counted daily for 8 days. Tetracycline-induced protein overexpression was tested by western blot analysis using monoclonal rat anti-HA antibodies. Values represent means ± SD of 3 independent replicates.

To evaluate the effect of overexpression of *Tc*BDF2HA and *Tc*BDF2HAdm on epimastigote replication we performed growth curves in the presence (tet+) and absence (tet-) of tetracycline for 8 days. As seen in Figures 3A and B (left panel), the expression of wild type and mutant *Tc*BDF2 affects epimastigote growth, when compared with the uninduced parasites. In fact, the effect seems to be higher when the wild type version is overexpressed. When we compared (ANOVA test, *p* > 0,05) the growth rate at day 8 was significatively different between the lines (34,5% for BDF2 and 49,3% for BDF2m, in both cases considering 100% the growth of corresponding non-induced cultures). This effect was observed in all the experiments we did with these lines. Moreover, the growth rate of the partially truncated mutant (*Tc*BDF2Δ177HA) and the single amino acid (*Tc*BDF2W92A) mutant had an intermediate effect when overexpressed, to the wild type and double mutant TcBDF2 (data not shown). These data suggest that the correct endogenous level of a functional *Tc*BDF2 is essential for the normal progression of the epimastigote cell cycle since overexpression of *Tc*BDF2HA or of a protein unable to recognize the acetylated lysine (*Tc*BDF2HAdm), even in low amounts produces a halt in epimastigote normal proliferation. However, the effect by which the wild type or mutant *Tc*BDF2 overexpression impairs parasites growth seems to be different. When the distribution of parasites along the cell cycle was tested by flow cytometry in non-synchronized cultures, no evident differences were observed between non-induced and *Tc*BDF2-induced cultures until 72 hs. In contrast, in *Tc*BDF2dm-induced cultures a population of parasites with DNA content lower than 2n appears, which was increased from approximately 5% at 24 hs to 8.2 % at 72 hs (Supplementary Figure 2). The appearances of this population could indicate that these parasites may be entering either an apoptotic or necrotic process. This phenomenon support the idea that the mechanisms by which overexpression of wild type or mutant *Tc*BDF2 affect the parasites are not the same.

### *Tc*BDF2 affects *T. cruzi* performance on mammalian host Infection

To better evaluate the potential of *Tc*BDF2 as a druggable target for Chagas disease treatment, we examined the involvement of this bromodomain factor on *T. cruzi* infective stages in the mammalian host. We evaluated if *Tc*BDF2 was able to modulate the infectivity of the parasite by allowing the parasites to invade Vero cells with *T*. *cruzi* transgenic lines overexpressing *Tc*BDF2HA or *Tc*BDF2HAdm. Given the advantage of the overexpression system which allows us to induce the production of the proteins at different time points, we were able to analyze how the overexpression of *Tc*BDF2HA or *Tc*BDF2HAdm affects *T. cruzi* performance throughout the entire infection process. Briefly, trypomastigotes were pre-incubated with tetracycline (Figure 4, +/- and +/+ condition) or not (-/+ and -/- condition) and then allowed to infect a monolayer of Vero cells. After 6 hs free trypomastigotes were washed out and fresh medium with (+/+ and -/+ condition) or without tetracycline (+/- and -/- condition) was added to the cells and incubated for 3 days. The -/- condition represents the non-induced control where tetracycline was never added to the medium and the +/+ condition shows the effect of protein overexpression during the whole infection assay.

**Figure 4:**
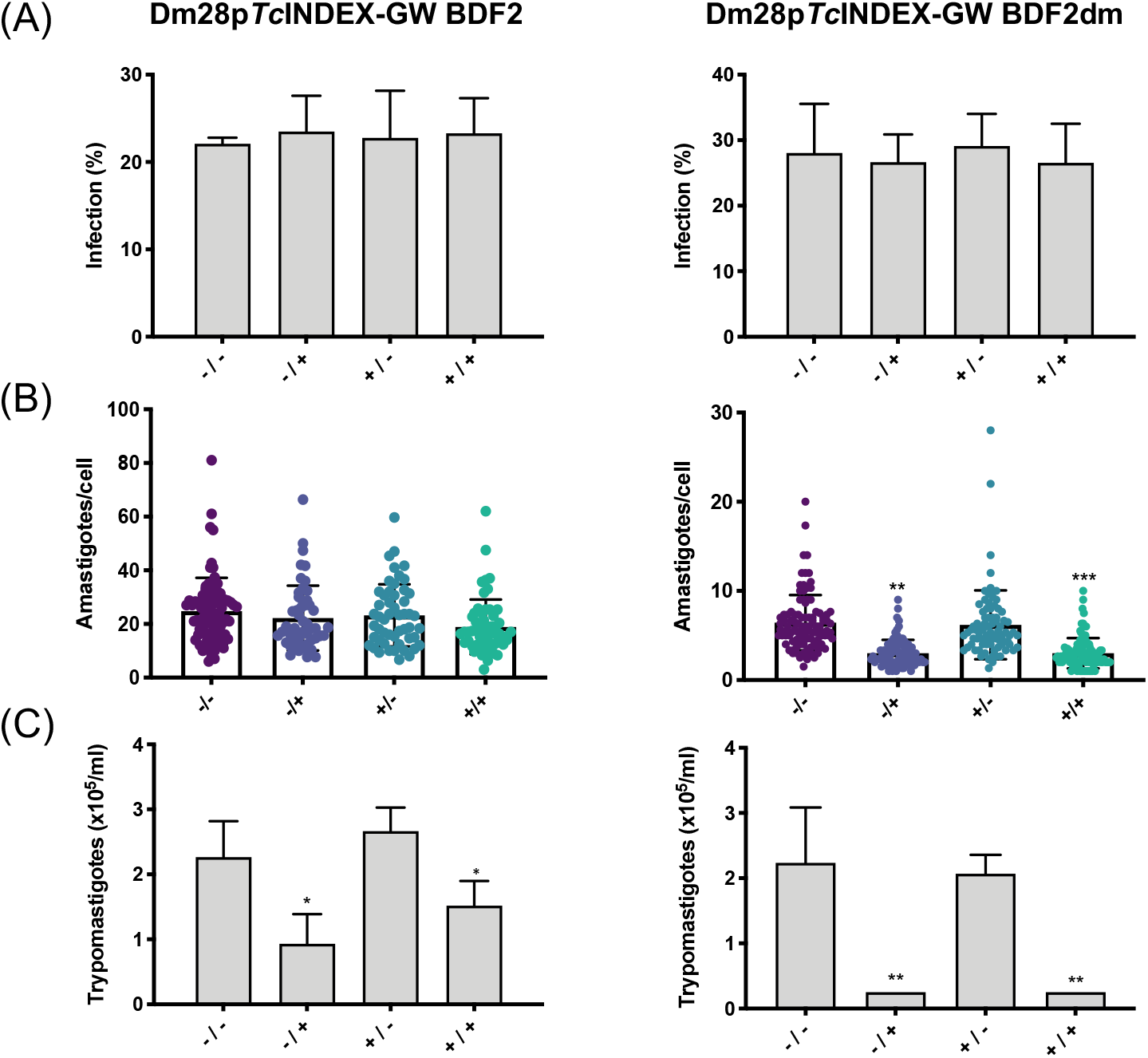
Overexpression of *Tc*BDF2HAdm alters intracellular replication and trypomastigote release. The infection performance of *T. cruzi* Dm28c transfected with p*Tc*INDEX-GW-*Tc*BDF2HA or p*Tc*INDEX-GW-BDF2HAdm were analyzed in the absence (Tet-) or presence (Tet+) of 0.5 μg/ml tetracycline. To test the effect over the different phases of infection, we designed different induction protocols as follows: (-/-), Tet was never added to the medium, noninduced control; (+/-), trypomastigotes were pre-treated with Tet for 2 hours before infection, and during the invasion incubation period but not after; (-/+), trypomastigotes were not induced, Tet was only added after infection to see the effect on the amastigote stage; (+/+), trypomastigotes were pre-treated and Tet was present during the whole infection assay, to see the overexpression effect on both trypomastigotes and amastigotes. The percentage of infected cells (A), the number of amastigotes per cell (B), and the number of trypomastigotes released 6 days post-infection (C) were determined by counting Giemsa-stained slides using a light microscope. Results are expressed as means ± SD of three independent biological experiments. Statistical analysis of the data was carried out using one-way ANOVA using the Multiple Comparison option, where the (-/-) condition was considered the control group, *\*p* < 0.05, *\*\*p* < 0.001, and *\*\*\*p* < 0.005.

First, we measured the ability of the overexpressing trypomastigotes to invade Vero cells. As can be seen in Figure 4A, the percentage of cells infected under the conditions (+/-) and (-/-) showed no statistical differences for both *Tc*BDF2HA and *Tc*BDF2HAdm. Trypomastigotes are considered the form with the lowest levels of active transcription in the life cycle of *T. cruzi* ^52^. Considering that *Tc*BDF2 is a protein that participates in complexes related to the transcriptional capacity of chromatin, it is not surprising that an imbalance of this protein, whether in its wild type or mutated form, does not have a marked effect on the physiology of this stage of the parasite.

Moving forward in the infection process we also evaluated how *Tc*BDF2 affects amastigote proliferation. Intracellular amastigote replication was measured as the average number of intracellular amastigotes per infected cell at 72 hs post-infection. Figure 4B represents the average numbers of amastigotes per cell for all the induction conditions when both *Tc*BDF2HA and *Tc*BDF2HAdm were overexpressed. As it can be observed, when *Tc*BDF2HA was overexpressed we did not detect any significant difference in the number of amastigotes among all the conditions assayed but strikingly we observed a significant decrease of about 50% in the amastigote proliferation when *Tc*BDF2HAdm was overexpressed. It is worth to highlight that this behavior was opposite to that observed in epimastigotes, where the overexpression of the wild type protein triggers the higher detrimental effect as shown in Figure 1.

Finally, we focused on the amastigote/trypomastigote differentiation and trypomastigote release from the infected cells. After 4 days post-invasion, trypomastigotes started to be released to the medium from the infected Vero cells. To analyze how *Tc*BDF2 impacts these final steps it is important to compare the (-/+) vs (-/-) conditions taking into account that the final result is a sum of all the processes happening once the initial trypomastigote invades the cell until the new trypomastigotes are released for further infections. As it can be seen in Figure 4C, the overexpression of *Tc*BDF2HA affects negatively the final steps reducing approximately by 50% the number of released parasites. However, when *Tc*BDF2HAdm was overexpressed we noticed a sharp decrease in trypomastigote released since parasites were practically undetectable in the culture medium.

Taken together, all these results suggest that *Tc*BDF2, or possibly the complexes that include it, are essential for epimastigote and amastigote replication and for amastigote differentiation to trypomastigotes. However, the dissimilar response of the different life cycle stages to the increased levels of *Tc*BDF2 and *Tc*BDF2dm suggests a complex landscape, where more than one protein complex could be playing a variety of roles. Even though the elucidation of the composition of these complexes and their function is beyond the objectives of this work, our results support *Tc*BDF2 as a target for the development of new drugs against Chagas disease.

### Recombinant *Tc*BDF2HA interacts with bromodomain inhibitors

Once demonstrated that *Tc*BDF2 is essential for almost all life cycle stages of *T. cruzi*, we tested the ability of small molecules that can act as bromodomain inhibitors to bind *Tc*BD2. In this case, we made use of an autofluorescence quenching assay, due to the presence of a tryptophan in the hydrophobic pocket, previously validated with *Tc*BDF3 ^42^. Briefly, we obtained the intrinsic fluorescence spectrum of each protein and evaluated the changes they underwent upon adding increasing amounts of each inhibitor (Figure 5). In this case, we assayed the recombinant bromodomain portion of *Tc*BDF2 (*Tc*BD2) and *Tc*BDF2HAdm (*Tc*BD2dm). Both fragments were obtained by cloning and overexpressing them in a soluble manner in *E.coli-BL21* using the commercial plasmid pDEST17. The folding of the purified bromodomains was compared by circular dichroism, corroborating that the spectra, corresponding to the presence of alpha helixes, was identical for both the wild type and the mutated domain (Supplementary Figure 3).

**Figure 5:**
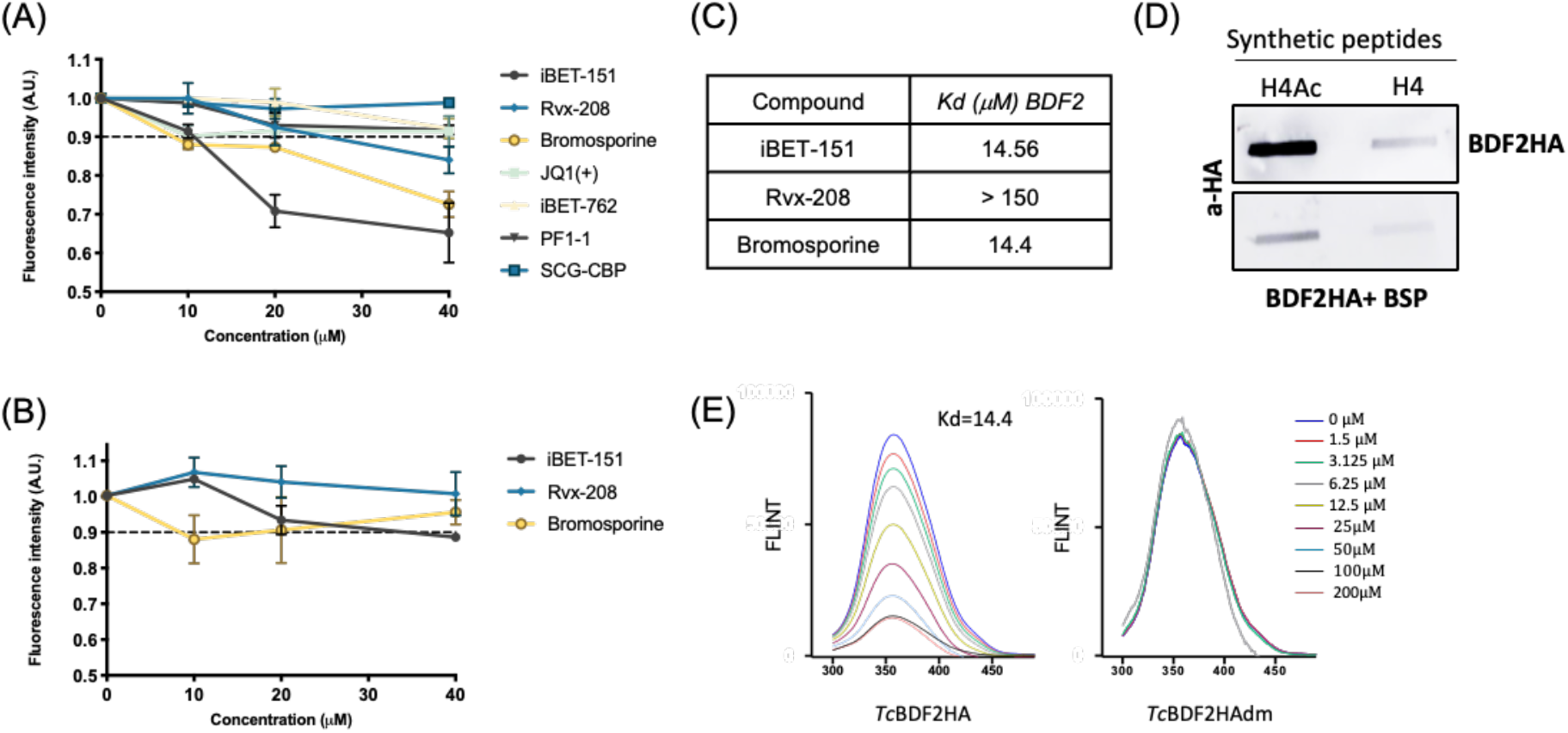
*Tc*BDF2HA interacts with acetylated histone H4 and three human bromodomain inhibitors. Normalized intrinsic fluorescence of recombinant *Tc*BDF2HA (A) and *Tc*BDF2dmHA (5 μM) (B) with an increasing amount of the indicated bromodomain inhibitor (10, 20, 40 μM). (C) Dissociation constants (*Kd*) of iBET-151, Rvx-208, and Bromosporine (BSP) were calculated from double logarithmic plots of the intrinsic fluorescence spectra of BDF2HA with the compounds. (D) Slot far-western blot assay. Tri-acetylated and non-acetylated (H4Ac) histone H4 (H4) peptides were blotted onto a nitrocellulose membrane and incubated with HA-tagged recombinant *Tc*BDF2 in the absence and presence of Bromosporine. The bound recombinant protein was detected with anti HA antibodies (aHA). (E) Fluorescence spectra of BDF2wt (left) and BDF2m (right) with increasing concentrations of BSP (0 to 200μM) and fixed concentration of protein 30 μM.

As can be observed in Figure 5A, *Tc*BD2 intrinsic fluorescence decreased in the presence of the human bromodomain inhibitors iBET-151, RVX-208, and Bromosporin (BSP) but remained unaltered with the other inhibitors assayed. However, no variation in the intrinsic fluorescence of the mutated version (Figure 5B) was observed when incubated with these compounds, indicating that the interaction depends on the integrity of the *Tc*BDF2 hydrophobic pocket. The data was further examined using the modified Stern-Volmer equation to calculate the dissociation constants (*Kd*), which was around 14 μM for iBET-151 and BSP but significantly higher for Rvx-208 (Figure 4C). *Kd* values we also calculated by thermal shift obtaining similar results. The complete fluorescence spectra of the wild type and mutant *Tc*BDF2 in the presence of increasing concentrations of BSP are shown in Figure 5E. BSP also blocked the interaction of recombinant *Tc*BDF2 with a peptide derived from histone H4 acetylated on lysines 4, 10, and 14, as seen by far-western blot (Figure 5D). This supports the idea that the inhibitors are recognized by the hydrophobic pocket of the bromodomain and compete with its ligand.

The affinity to *Tc*BDF2 obtained for iBET-151 and RVX-208 was at least two orders of magnitude lower than those reported for their specific targets in mammalian cells. iBET151 displays potent binding affinity to human BETs (BRD2, BRD3, BRD4, and BRD9) with *Kd* of 0,02 μM to 0,1 μM, but does not bind at least 23 other bromodomain proteins. It blocks the recruitment of BET to chromatin *in vivo* and has shown therapeutic activity in distinct mouse models of murine and human leukemias ^53^. RVX-208 (also known as Apabetalon) is an inhibitor with selectivity for the second BD of BET transcriptional regulators (*Kd* = 0.51 μM) ^54^, Affinity was also low for BSP, but in contrary to iBET-151 and RVX-208, this is a broad-spectrum inhibitor for bromodomains and considered a very useful tool for elucidating biological roles of reader domains as well as for the validation of functional assays ^55^. Also, *Tc*BDF2 was co-crystallized previously with BSP at its hydrophobic pocket. When we modeled by local docking this interaction using RosettaLigand ^56^, BSP was found at the same position as in the crystal structure indicating that our settings were correct (Supplementary Figure 4A-B). Using the same approach we modeled the interaction of *Tc*BDF2 with the inhibitors we tested *in vitro* and calculated the interphase energy and the root mean square deviation (RMSD) (Supplementary Figure 4C). The results of this *in silico* evaluation of the interactions between *Tc*BDF2 and the compounds mentioned in Figure 5 correlates with the dissociation constants obtained by quenching of the intrinsic tryptophan fluorescence, further validating this assay (Supplementary Figure 4D).

When our results are compared with those obtained previously by analyzing the same panel of bromodomain inhibitors against recombinant *Tc*BDF3 ^57^, it can be clearly seen that *Tc*BDF2 and *Tc*BDF3 have different binding specificities. iBET-151 binds to both bromodomains with a similar affinity, but JQ1(+), which binds *Tc*BDF3 with an affinity similar to iBET-151, does not interact with *Tc*BDF2. In contrast, BSP shows a higher affinity for *Tc*BDF2 than for *Tc*BDF3. On the other hand, there is no direct correlation between the affinity of the inhibitors we assayed with both bromodomains and their activity against the parasite. We previously showed that iBET-151 and JQ1(+) have the lower IC_50_ against Dm28c epimastigotes (6,35 μM and 7.14 μM respectively). The *Kd* estimated of iBET-151 was similar for *Tc*BDF2 and *Tc*BDF3. In contrast, BSP has a *Kd* for *Tc*BDF2 very similar to that of iBET-151, but low activity against epimastigotes (IC_50_ > 50 μM). This discrepancies between *in vitro* measured *Kd* and *in vivo* obtained IC_50_ are not surprising and can be assigned to many factors, including accessibility of the compounds to cell nucleus. Unfortunately, due to the toxicity of this panel of compounds over mammalian transformed cell lines that are normally used for infection experiments, like Vero or HeLa, their activity against amastigotes cannot be determined.

As it was already mentioned, some bromodomains inhibitors were also previously tested against *T. brucei.* The affinity ranges obtained were lower than those described herein. In addition, results are significantly different when the orthologs are compared. Shultz *et al.* ^46^ showed that iBET-151 binds to *Tb*BDF2 and *Tb*BDF3 with a a *Kd* of 225 μM and 175 μM respectively. Surprisingly, when iBET-151 was co-crystalized with *Tb*BDF2 it was found in a completely atypical position, flipped by roughly 180° to the position it binds to human BDs, something that could explain the low affinity of the interaction. Later, Yang *et al*. described a compound, GSK2801, that binds with high affinity (*Kd* = 15 μM) to the BD from *Tb*BDF2 and with low affinity (*Kd* = 83 μM) to the second BD from *Tb*BDF5. In contrast, GSK2801 did not bind to *Tb*BDF3 and neither to the first BD from *Tb*BDF5 ^16^.

## Conclusions

We have previously suggested that *Tc*BDF1 and *Tc*BDF3 are two essential bromodomain-containing proteins for the parasite to accomplish their life cycle by using a dominant-negative approach. The fact that it was not possible to generate full knockouts for these genes by CRISPR/Cas9 support this hypothesis (data not published). Herein, using similar tools, we showed that *Tc*BDF2 is essential for both epimastigote and amastigote growth. Determination of essentiality of genes in *T. cruzi* is a problem associated with the lack of appropriated genetic tools, like inducible knock-outs or knock-downs. Even though the availability of efficient CRISPR/Cas9 protocols overcame the historical difficulties to generate knock-outs by homologous recombination, the plasticity and tendency to recombine of the *T. cruzi* genome still makes it a hard task. The repeated failure of producing a complete knock-out by any of these techniques is a strong indication of essentiality but certainly gives no certitude. We complement the knock-out approach with a dominant-negative experimental setup, based on the assumption that BDFs take part in multiprotein complexes, in which they carry the binding activity to the acetylated-target protein. Then, mutations in the amino acids essential to this binding activity will be incorporated into de complexes and transform them into non-functional, and giving an observable phenotype. Since in this approach the wild-type protein remains present, some functional complexes are also present and the strongness of the phenotype depends on the ratio between the wild-type and the mutated forms of the protein. In our case, we used the high-expression inducible plasmid p*Tc*INDEX and we chose to mutate the amino acids Y85 and W92 in order to avoid acetyl-lysine binding but assure that the rest of *Tc*BDF2 is correctly folded as not to disturb its correct interaction with the partners in the complex. We tested both desired characteristics, correct folding and lack of binding, using the recombinant purified proteins. Under these conditions, the effect of *Tc*BDF2 can be considered a dominant-negative mutant. In the case of *Tc*BDF2, the overexpression of the wild-type protein also gives a phenotype indicating that large amounts of functional protein also disturb the function of the complexes. Taking into account that some components of the nuclear complexes that contain BDFs are promiscuous and participate in more than one complex we can imagine that overexpression of *Tc*BDF2 can disturb these complexes by changing the stoichiometry of their components in the nucleus. Another possibility to explain this effect is that free *Tc*BDF2 could bind acetylated histones and block them making them unable to bind to functional complexes. Even though we still lack evidence to support these speculations, both hypotheses could explain why the effect of *Tc*BDF2 overexpression is different from that of the dominant-negative mutant. In any case, all these experimental observations taken together, give us certitude about the essentiality of the function of *Tc*BDF2.

We also showed that bromodomain inhibitors can bind to *Tc*BDF2 through its hydrophobic pocket. However, the specific binding profile is different for *Tc*BDF2 than for *Tc*BDF3. The affinities of the tested inhibitors for *Tc*BDF2 are not high enough to be seriously considered lead compounds. In fact, we used them as probes to assay the concept of BD inhibitors as potential trypanocidal drugs. Our results, together with those obtained for other parasites, show that parasitic bromodomains, despite their structural preservation, are a family with a great substrate specificity diversity. This is observed both between proteins from different species (like *T. brucei* and *T. cruzi*) and between proteins from the same species, as we observed for *Tc*BDF2 and *Tc*BDF3. Although it is hard to imagine the existence of one compound that could inhibit all *T. cruzi* BDs without crossing with human BDs, results presented here and in the bibliography establish that it could be possible to find molecules with high binding specificity that can discriminate between one or more than one parasitic bromodomain and human bromodomains. Further studies about BDF function are certainly needed to better define a strategy for the search for new antiparasitic compounds based on this idea.

## Methods

### Parasite cultures

*T. cruzi* Dm28c epimastigotes were cultured at 28 °C in LIT medium (5 g/L liver infusion, 5 g/L bacto-tryptose, 68 mM NaCl, 5.3 mM KCl, 22 mM Na2HPO4, 0.2% (w/v) glucose and 0.002% (w/v) hemin) supplemented with 10% (v/v) heat-inactivated, UV-irradiated Fetal Calf Serum (FCS) (Internegocios S.A, Argentina).

### Molecular cloning of *Tc*BDF2HA

*Tc*BDF2 gene (*TC*DM_14496) was amplified by PCR from *T. cruzi* Dm28c genomic DNA using the following oligonucleotides: *Tc*BDF2HAw, designed to add a hemagglutinin (HA) tag at the N-terminus of the protein (5’-AAA<u>GGATCC</u>ATGTATCCGTATGATGTGCCGGATTATGCTGGGAAGCGTGGGCGT-3’), and *Tc*BDF2Rv (5’-AAA<u>GATATC</u>TTATTCATCATCACTCTCATCATCATAATAAAACTCTTC-3’). Truncated protein was constructed using the following reverse oligonucleotides: ΔC177-*Tc*BDF2Rv (5’-AAA<u>GATATC</u>TTAATTCTTGCCTCCTTTCTTCTTTTCCACAC-3’), ΔC162-*Tc*BDF2Rv (5’-AAA<u>GATATC</u>TTAAAGCTCCTCCAG-3’) ΔC117-*Tc*BDF2Rv (5’-AAA<u>GATATC</u>TTACGCGTCGTCGTCAACACTCAG-3’). The last construction corresponds to the bromodomain portion of *Tc*BDF2 and was employed to purify it as a recombinant protein for in *vitro* assays. The double mutant *Tc*BDF2HAdm (*Tc*BDF2Y85A/W92A) was constructed using a PCR-based site-directed mutagenesis strategy with the following oligonucleotides: *Tc*BDF2Y85AFw (5’-CTGCGAGAAGGCTAACGGCG-3’), *Tc*BDF2Y85ARv (5’-CGCCGTTAGCCTTCTCGCAG-3’), *Tc*BDF2W92AFw (5’CGACTCCGCTGCGGTTAAAG-3’) and *Tc*BDF2W92ARv (5’-CTTTAACCGCAGCGGAGTCG-3’).The PCR products were first cloned into the pCR2.1-TOPO vector (Invitrogen) and sequenced. The coding sequences were subcloned into a pENTR-3C vector (Invitrogen) using BamHI/EcoRV restriction sites included in the oligonucleotides (underlined) and then transferred to the p*Tc*INDEX-GW and to pDEST™17 (Gateway System, Invitrogen) vector by recombination using LR clonase II enzyme mix (Invitrogen).

### *Tc*BDF2 KO line generation

The general strategy was carried out as is indicated by Lander *et. al*^58^. Briefly, the SgRNA carrying the specific protospacer sequence was amplified from pUC_sgRNA plasmid using the oligonucleotides SgRNAFw (5’-GATC<u>GGATCC</u>GCGTGAGGGGACTTTGGACAGTTTTAGAGCTAGAAATAGC-3’) and SgRNARv (5’-CAGT<u>GGATCC</u>AAAAAAGCACCGACTCGGTG-3’) and cloned into Cas9/pTREX-n plasmid by using BamHI site (underlined in the primers). Donor DNA was generated by PCR using as a template the blasticidin-S deaminase gene (*Bsd*). Ultramers utilized were *Tc*Bdf2KOFw (5’-ATAAAAGAAGAGGCACACAGAAAACAAAGAAAGCCGTCGAAATAATAAAGAAACAACAACAACAACAAGAGAGGCCTGGAGGTAACAGTATCGGGGGGTGATGGCCAAGCCTTTGTCTCA-3’) and *Tc*Bdf2KORv (5’-GCTGTCGTACTTTCCGGACTCAATGCCCTGACGAATTGTCGACAGGTCAACAGGATAATTTATGACCTTGTGGTAGTCCGGCAGCTCCGCAGCGCTGATTTTAGCCCTCCCACACATAAC-3’). The sequences of the 120 pb ultramers included the homology region for DNA recombination. *T. cruzi* DM28c epimastigotes were co-trasfected with the donor DNA and Cas9/pTREX-n plasmid and the parasites were selected by adding G418 and blasticidin in the medium as indicated in ^58^. A scrambled sgRNA (Scr-sgRNA/Cas9/pTREX-n) was used as control as described by Lander *et al*^58^.

### Protein purification

Protein purification pDEST17-*Tc*BDF2 and pDEST17-*Tc*BDF2m were transformed into *Escherichia coli BL21,* and the recombinant proteins (fused to a His-tag) were obtained by induction with 0.1 mM isopropyl-b-D-thiogalactopyranoside overnight at 22 °C. The protein was purified using Ni-NTA (Thermo Fisher™) following the manufacturer’s instructions. Purified proteins were dialyzed against 0.1 M phosphate buffer pH 8. The secondary structure of soluble proteins was measured by circular dichroism spectroscopy using a spectropolarimeter (Jasco J-810, Easton, MD, USA).

### Transgenic parasite generation

Epimastigotes from *T. cruzi* Dm28c were transfected with the pLEW13 plasmid to generate parasites expressing T7 RNA polymerase and the Tet repressor using a standard electroporation method. Briefly, epimastigotes were cultured in LIT medium at 28 °C to a final concentration of 3-5 × 10^7^ parasites/ml. Then, parasites were harvested by centrifugation at 1500 g for 5 min at room temperature, washed twice with phosphate-buffered saline (PBS), and resuspended in 0.35 ml transfection buffer (0.5 mM MgCl2, 0.1 mM CaCl2 in PBS, pH 7.5) to a density of 1 × 10^8^ cells/ml for each transfection. Electroporation was performed in a 0.2 cm gap cuvette (Bio-Rad) with ~40 μg of plasmid DNA added to a final volume of 400 μl. The parasite-DNA mixture was kept on ice for 25 min and subjected to a 450 V, 500 μF pulse using GenePulser II (Bio-Rad, Hercules, USA). After electroporation, cells were transferred into 3 ml of LIT medium containing 10% FCS, maintained at room temperature for 15 minutes, and then incubated at 28 °C. Geneticin (G418; Life Technologies) was added at a concentration of 200 μg/ml, and parasites were incubated at 28 °C. After selection, pLEW13 transfected epimastigotes were maintained in the presence of 200 μg/ml of G418. This parental cell line was then transfected with p*Tc*INDEX-GW-*Tc*BDF2HA (WT and mutant versions) plasmids following a similar protocol and transgenic parasites were obtained after 3 weeks of selection with 100 μg/ml G418 and 200 μg/ml Hygromycin B (Sigma).

### *T. cruzi* infection of Vero cells

Vero cells (ATCC CCL-81) were cultured in Dulbecco’s Modified Eagle Medium (DMEM) (ThermoFisher), supplemented with 2 mM L-glutamine, 10% FCS. For the first round of infection, metacyclic trypomastigotes were obtained by spontaneous differentiation from late-stationary phase cultured epimastigotes at 28 °C. Cell-derived trypomastigotes were obtained by infection with metacyclic trypomastigotes in Vero cell monolayers. For the infection and amastigote proliferation experiments, we used cell-derived trypomastigotes released from the second round of infection. Trypomastigotes were collected from the supernatant of the infected cells culture, harvested by centrifugation at 5000 g for 10 min at room temperature, resuspended in DMEM, and counted in a Neubauer chamber. When indicated, the parasites were pre-incubated with Tet (0.5 μg/ml) for 2 hs to allow protein induction, and a new monolayer of cultured Vero cells was infected with a MOI of 10:1. After a 6 hs-incubation at 37 °C, the free trypomastigotes were removed by washes with PBS. This pre-treatment of trypomastigotes with tetracycline before infection and during the 6 hs-incubation to allow invasion is indicated with a “+” before the bar “(+/)”. Then, infected Vero cell cultures were incubated in DMEM supplemented with 2% FCS with or without Tet (0.5 μg/ml) for three days. The presence of Tet during this incubation period when amastigotes replicate inside the infected cell is indicated by the “+” after the bar “(/+)”. The experiments were stopped by cell fixation with methanol and the percentage of infected cells and the mean number of amastigotes per infected cell were determined by direct slide counting (Nikon Eclipse Ni-U microscope). Giemsa staining was used for amastigotes visualization and approximately 800 cells were counted per slide. Alternatively, after 4 days post-infection, the supernatant from monolayer infection was taken to determine the number of trypomastigotes released from cells. The significance of the results was analyzed with two-way ANOVA using GraphPad PRISM version 6.0 for Mac (GraphPad Software, La Jolla, CA, USA). Results are expressed as means ± SEM of triplicates, and represent one of three independent experiments performed.

### Western blot analysis

Protein extracts were fractioned in SDS-PAGE and transferred to a nitrocellulose membrane. Transferred proteins were visualized with Ponceau S staining. Membranes were treated with 10% non-fat milk in PBS for 2 hs and then incubated with specific antibodies diluted in 0.5% Tween20 in PBS (PBS-T) for 3 hs. Antibodies used were: rat monoclonal anti-HA (ROCHE), affinity-purified rabbit polyclonal anti-BDF2, mouse monoclonal anti-trypanosome α-tubulin clone TAT-1 (a gift from K. Gull, University of Oxford, UK). Bound antibodies were detected using peroxidase-labeled anti-rabbit IgG (GE Healthcare), or anti-rat IgG (Thermo Scientific) and developed using ECL Prime kit (GE Healthcare) according to manufacturer’s protocols. Immunoreactive bands were visualized using an Amersham™ Imager 600 digital imager.

### Immunofluorescence

Mid-log epimastigotes were harvested by centrifugation at 1500 g for 5 min at room temperature and washed twice before fixation in 4% paraformaldehyde solution. Fixed parasites were placed on a coverslip pre-coated with poly L-lysine for 20 min and then washed with PBS. Permeabilization was done with 0.2% Triton X-100 solution for 10 min. After washing with PBS, parasites were incubated with the appropriate primary antibody diluted in 1% BSA in PBS for 2 hs at room temperature. Non-bound antibodies were washed with PBS-T and then the slides were incubated with anti-HA (Roche) and anti-rat IgG::FITC (Life Technologies) and 2 μg/ml 4’,6-diamidino-2-fenilindol (DAPI) for 1 h. The slides were washed again with PBS-T and mounted with VectaShield (Vector Laboratories). Images were acquired with a confocal microscope ZEISS LSM 880. Adobe Photoshop CS and ImageJ software were used to process all images.

### Flow cytometry

One million cells were fixed with cold 70% ethanol and then washed with PBS and stained with 20 μg/ml Propidium Iodide (PI) in buffer K (0.1% sodium citrate, 0.02 mg/ml RNAse A (Sigma), and 0.3% NP-40. Ten thousand events per sample were acquired using BD Cell Sorter BD FACSAria II. Results were analyzed using FlowJo software.

### Far Western blot and slot blot

Far Western blots were performed using 10 μg of each 16 amino acid-long peptides derived from the histone H4 sequence, or the same sequence with acetylated lysines 4, 10, and 14 (CAKGKacKSGEAKaGTQKaRQ, immobilized onto a nitrocellulose membrane. Peptides were visualized by Ponceau S staining. The membranes were treated with 5% non-fat milk in PBS (blocking buffer) for 2 hs, and then with recombinant *Tc*BDF2HA diluted in blocking buffer (0.1 and 0.5 μg/ml) for 1 h. After this incubation, membranes were washed with PBS Tween 0.1%. Bound proteins were visualized using rat anti-HA antibody and detected using peroxidase-labeled anti-rat IgG antibody and ECL Plus (GE Healthcare).

### Protein purification

pDEST17-*Tc*BDF2HA and pDEST17-*Tc*BDF2HAdm were transformed into *Escherichia coli* BL21, and the expression of recombinant proteins (fused to a His-tag and haemagglutinin tag) were induced with 0.1 mM isopropyl-β-D-thiogalactopyranoside during 5 hs at 37 °C. The proteins were purified under native conditions by affinity chromatography using Ni–NTA agarose (Qiagen) following the manufacturer’s instructions.

### Fluorescence spectroscopy

A 2 mL solution containing 5 μM recombinant *Tc*BDF2HA or *Tc*BDF2HAdm was titrated by successive addition of the bromodomain inhibitors in 96-well microplates (Fluotrac 200 black, GreinerBioOne). All determinations were made from the top with an excitation wavelength of 275 nm and a 340/30 nm emission filter, 50 reads per well, and a PMT-sensitivity setting of 180 in a Synergy HT multidetector microplate reader equipped with time-resolved capable optics. For BSP (0 to 200 μM) fluorescence spectra were acquired with an excitation wavelength of 295 nm and emission was recorded in the range of 300–450 nm and 30 μM recombinant *Tc*BDF2HA or *Tc*BDF2Hadm. All fluorescence measurements were corrected with a blank solution and with the emission spectra of each concentration of inhibitor. Taking into account the inner filter effect in the quenching process, we corrected the fluorescence intensity of *Tc*BDF2 using the following equation ^59^: F_corr_ = F_obs_X 10^(Aexc+Aem)/2^, where F_corr_ and Fobs are the corrected and observed fluorescence intensity of *Tc*BDF2HA, and A_exc_ and A_em_ are the absorption values of the system at the excitation and emission wavelength, respectively.

### Statistical analysis

Experiments were performed in triplicate, and at least three independent experiments were performed. Data are presented as the mean ± SEM. Statistical analysis of the data was carried out using two-way ANOVA and unpaired two-tailed Student’s t-test. Differences between the experimental groups were considered significant as follows: *\*P* < 0.05, *\*\*P* < 0.001 and *\*\*\*P* < 0.005.

## Acknowledgments

This work was supported by Agencia Nacional de Ciencia y Tecnologia from Argentina (PICT-2014-2593, PICT-2017-1978), Universidad Nacional de Rosario (1BIO490), Research Council United Kingdom (MR/P027989/1) and GlaxoSmithKline. We would like to thank Dolores Campos and Dr. Romina Manarin for their technical assistance with Vero cell and parasites culture, Mara Ojeda for technical assistance with flow cytometry, and Dr. Teresa Cruz-Bustos for discussions on the use of CRISPR/Cas9. Part of the results presented in this work have been obtained by using the facilities of the CCT-Rosario Computational Center, member of the High Performance Computing National System (SNCAD, MincyT-Argentina).

## Supporting Information

Expression of *Tc*BDF2dm in epimastigotes, cell cycle measurements by flow cytometry, Circular dichroism of recombinant *Tc*BDF2 and m*Tc*BDF2dm, validation of RosettaLigand scripts and correlation with dissociation constants.

## For Table of Contents Use Only

*Tc*BDF2 was characterized to determine its essentiality and potential as drug target against Chagas disease.

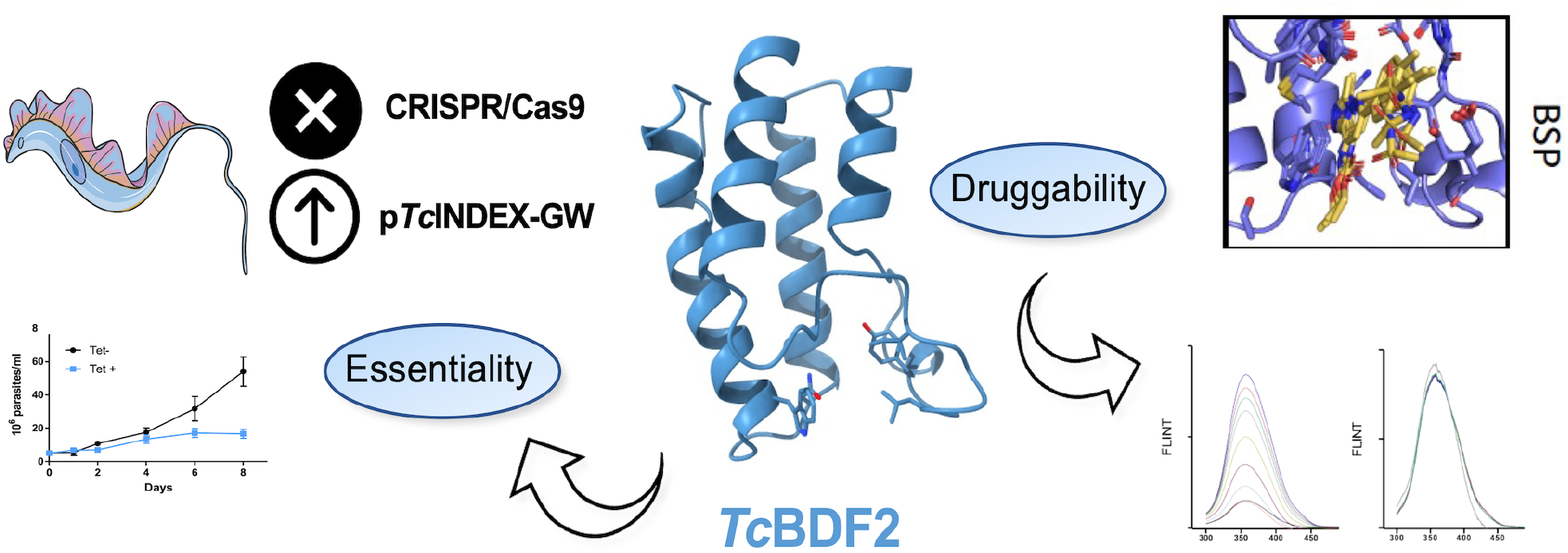

## References

(1) Requena-Méndez, A.; Aldasoro, E.; de Lazzari, E.; Sicuri, E.; Brown, M.; Moore, D. A. J.; Gascon, J.; Muñoz, J. Prevalence of Chagas Disease in Latin-American Migrants Living in Europe: A Systematic Review and Meta-Analysis. PLoS Negl. Trop. Dis. 2015. https://doi.org/10.1371/journal.pntd.0003540.

(2) WHO. WHO | Chagas disease (American trypanosomiasis) Factsheet.

(3) Sales, P. A.; Molina, I.; Murta, S. M. F.; Sánchez-Montalvá, A.; Salvador, F.; Corrêa-Oliveira, R.; Carneiro, C. M. Experimental and Clinical Treatment of Chagas Disease: A Review. American Journal of Tropical Medicine and Hygiene. 2017. https://doi.org/10.4269/ajtmh.16-0761.

(4) Martínez-Calvillo, S.; Vizuet-De-Rueda, J. C.; Florencio-Martínez, L. E.; Manning-Cela, R. G.; Figueroa-Angulo, E. E. Gene Expression in Trypanosomatid Parasites. Journal of Biomedicine and Biotechnology. 2010. https://doi.org/10.1155/2010/525241.

(5) De Gaudenzi, J. G.; Carmona, S. J.; Aġuero, F.; Frasch, A. C. Genome-Wide Analysis of 3’-Untranslated Regions Supports the Existence of Post-Transcriptional Regulons Controlling Gene Expression in Trypanosomes. PeerJ 2013. https://doi.org/10.7717/peerj.118.

(6) Clayton, C. E. Gene Expression in Kinetoplastids. Curr. Opin. Microbiol. 2016, 32, 46–51. https://doi.org/10.1016/j.mib.2016.04.018.

(7) Rudenko, G. Epigenetics and Transcriptional Control in African Trypanosomes. Essays Biochem. 2010. https://doi.org/10.1042/BSE0480201.

(8) Staneva, D. P.; Carloni, R.; Auchynnikava, T.; Tong, P.; Rappsilber, J.; Arockia Jeyaprakash, A.; Matthews, K. R.; Allshire, R. C. A Systematic Analysis of Trypanosoma Brucei Chromatin Factors Identifies Novel Protein Interaction Networks Associated with Sites of Transcription Initiation and Termination. Genome Res. 2021. https://doi.org/10.1101/gr.275368.121.

(9) Jones, N. G.; Geoghegan, V.; Moore, G.; Carnielli, J. B. T.; Newling, K.; Calderón, F.; Gabarró, R.; Martín, J.; Prinjha, R.; Rioja, I.; Wilkinson, A. J.; Mottram, J. C. Bromodomain Factor 5 Is an Essential Transcriptional Regulator of the Leishmania Genome. bioRxiv 2021, 2021.09.29.462384. https://doi.org/10.1101/2021.09.29.462384.

(10) Felsenfeld, G.; Groudine, M. Controlling the Double Helix. Nature. 2003. https://doi.org/10.1038/nature01411.

(11) Kouzarides, T. Chromatin Modifications and Their Function. Cell. 2007. https://doi.org/10.1016/j.cell.2007.02.005.

(12) Respuela, P.; Ferella, M.; Rada-Iglesias, A.; Åslund, L. Histone Acetylation and Methylation at Sites Initiating Divergent Polycistronic Transcription in Trypanosoma Cruzi. J. Biol. Chem. 2008. https://doi.org/10.1074/jbc.M802081200.

(13) Siegel, T. N.; Hekstra, D. R.; Kemp, L. E.; Figueiredo, L. M.; Lowell, J. E.; Fenyo, D.; Wang, X.; Dewell, S.; Cross, G. A. M. Four Histone Variants Mark the Boundaries of Polycistronic Transcription Units in Trypanosoma Brucei. Genes Dev. 2009. https://doi.org/10.1101/gad.1790409.

(14) Thomas, S.; Green, A.; Sturm, N. R.; Campbell, D. A.; Myler, P. J. Histone Acetylations Mark Origins of Polycistronic Transcription in Leishmania Major. BMC Genomics 2009. https://doi.org/10.1186/1471-2164-10-152.

(15) Reynolds, D.; Hofmeister, B. T.; Cliffe, L.; Alabady, M.; Siegel, T. N.; Schmitz, R. J.; Sabatini, R. Histone H3 Variant Regulates RNA Polymerase II Transcription Termination and Dual Strand Transcription of SiRNA Loci in Trypanosoma Brucei. PLoS Genet. 2016. https://doi.org/10.1371/journal.pgen.1005758.

(16) Yang, X.; Wu, X.; Zhang, J.; Zhang, X.; Xu, C.; Liao, S.; Tu, X. Recognition of Hyperacetylated N-Terminus of H2AZ by TbBDF2 from Trypanosoma Brucei. Biochem. J. 2017. https://doi.org/10.1042/BCJ20170619.

(17) Wedel, C.; Förstner, K. U.; Derr, R.; Siegel, T. N. GT -rich Promoters Can Drive RNA Pol II Transcription and Deposition of H2A.Z in African Trypanosomes. EMBO J. 2017. https://doi.org/10.15252/embj.201695323.

(18) Anderson, B. A.; Wong, I. L. K.; Baugh, L.; Ramasamy, G.; Myler, P. J.; Beverley, S. M. Kinetoplastid-Specific Histone Variant Functions Are Conserved in Leishmania Major. Mol. Biochem. Parasitol. 2013. https://doi.org/10.1016/j.molbiopara.2013.09.005.

(19) Schulz, D.; Zaringhalam, M.; Papavasiliou, F. N.; Kim, H.-S. Base J and H3.V Regulate Transcriptional Termination in Trypanosoma Brucei. PLoS Genet. 2016, 12 (1), e1005762. https://doi.org/10.1371/journal.pgen.1005762.

(20) Kraus, A. J.; Vanselow, J. T.; Lamer, S.; Brink, B. G.; Schlosser, A.; Siegel, T. N. Distinct Roles for H4 and H2A.Z Acetylation in RNA Transcription in African Trypanosomes. Nat. Commun. 2020. https://doi.org/10.1038/s41467-020-15274-0.

(21) Rosón, J. N.; Vitarelli, M. de O.; Costa-Silva, H. M.; Pereira, K. S.; Pires, D. da S.; Lopes, L. de S.; Cordeiro, B.; Kraus, A. J.; Cruz, K. N. T.; Calderano, S. G.; Fragoso, S. P.; Siegel, T. N.; Elias, M. C.; da Cunha, J. P. C. H2B.V Demarcates Divergent Strand-Switch Regions, Some TDNA Loci, and Genome Compartments in Trypanosoma Cruzi and Affects Parasite Differentiation and Host Cell Invasion. PLoS Pathog. 2022, 18 (2), e1009694. https://doi.org/10.1371/journal.ppat.1009694.

(22) Berná, L.; Rodriguez, M.; Chiribao, M. L.; Parodi-Talice, A.; Pita, S.; Rijo, G.; Alvarez-Valin, F.; Robello, C. Expanding an Expanded Genome: Long-Read Sequencing of Trypanosoma Cruzi. Microb. genomics 2018. https://doi.org/10.1099/mgen.0.000177.

(23) Zeng, L.; Zhou, M. M. Bromodomain: An Acetyl-Lysine Binding Domain. FEBS Lett. 2002, 513 (1), 124–128. https://doi.org/10.1016/s0014-5793(01)03309-9.

(24) Marmorstein, R.; Zhou, M.-M. Writers and Readers of Histone Acetylation: Structure, Mechanism, and Inhibition. Cold Spring Harb. Perspect. Biol. 2014, 6 (7), a018762. https://doi.org/10.1101/cshperspect.a018762.

(25) Zaware, N.; Zhou, M. M. Bromodomain Biology and Drug Discovery. Nature Structural and Molecular Biology. 2019. https://doi.org/10.1038/s41594-019-0309-8.

(26) Vidler, L. R.; Brown, N.; Knapp, S.; Hoelder, S. Druggability Analysis and Structural Classification of Bromodomain Acetyl-Lysine Binding Sites. J. Med. Chem. 2012. https://doi.org/10.1021/jm300346w.

(27) Sanchez, R.; Meslamani, J.; Zhou, M. M. The Bromodomain: From Epigenome Reader to Druggable Target. Biochimica et Biophysica Acta - Gene Regulatory Mechanisms. 2014. https://doi.org/10.1016/j.bbagrm.2014.03.011.

(28) Kulikowski, E.; Rakai, B. D.; Wong, N. C. W. Inhibitors of Bromodomain and Extra-Terminal Proteins for Treating Multiple Human Diseases. Medicinal Research Reviews. 2021. https://doi.org/10.1002/med.21730.

(29) Clinical Trials database. U.S. National Library of Medicine. NIH. 23 https://clinicaltrials.gov/.

(30) Alonso, V. L.; Ritagliati, C.; Cribb, P.; Cricco, J. A.; Serra, E. C. Overexpression of Bromodomain Factor 3 in Trypanosoma Cruzi (TcBDF3) Affects Differentiation of the Parasite and Protects It against Bromodomain Inhibitors. FEBS J. 2016. https://doi.org/10.1111/febs.13719.

(31) García, P.; Alonso, V. L.; Serra, E.; Escalante, A. M.; Furlan, R. L. E. Discovery of a Biologically Active Bromodomain Inhibitor by Target-Directed Dynamic Combinatorial Chemistry. ACS Med. Chem. Lett. 2018. https://doi.org/10.1021/acsmedchemlett.8b00247.

(32) Ramallo, I. A.; Alonso, V. L.; Rua, F.; Serra, E.; Furlan, R. L. E. A Bioactive Trypanosoma Cruzi Bromodomain Inhibitor from Chemically Engineered Extracts. ACS Comb. Sci. 2018. https://doi.org/10.1021/acscombsci.7b00172.

(33) Chua, M. J.; Robaa, D.; Skinner-Adams, T. S.; Sippl, W.; Andrews, K. T. Activity of Bromodomain Protein Inhibitors/Binders against Asexual-Stage Plasmodium Falciparum Parasites. Int. J. Parasitol. Drugs Drug Resist. 2018. https://doi.org/10.1016/j.ijpddr.2018.03.001.

(34) Hanquier, J.; Gimeno, T.; Jeffers, V.; Sullivan, W. J. Evaluating the GCN5b Bromodomain as a Novel Therapeutic Target against the Parasite Toxoplasma Gondii. Exp. Parasitol. 2020. https://doi.org/10.1016/j.exppara.2020.107868.

(35) Brand, M.; Clayton, J.; Moroglu, M.; Schiedel, M.; Picaud, S.; Bluck, J. P.; Skwarska, A.; Chan, A. K. N.; Laurin, C. M. C.; Scorah, A. R.; Larissa See, K. F.; Rooney, T. P. C.; Fedorov, O.; Perell, G.; Kalra, P.; Cortopassi, W. A.; Christensen, K. E.; Cooper, R. I.; Paton, R. S.; Pomerantz, W. C. K.; Biggin, P. C.; Hammond, E. M.; Filippakopoulos, P.; Conway, S. J. Controlling Intramolecular Interactions in the Design of Selective, High-Affinity, Ligands for the CREBBP Bromodomain. ChemRxiv 2020. https://doi.org/10.26434/chemrxiv.12081999.

(36) Tallant, C.; Bamborough, P.; Chung, C. W.; Gamo, F. J.; Kirkpatrick, R.; Larminie, C.; Martín, J.; Prinjha, R.; Rioja, I.; Simola, D. F.; Gabarró, R.; Calderón, F. Expanding Bromodomain Targeting into Neglected Parasitic Diseases. ACS Infect. Dis. 2021. https://doi.org/10.1021/acsinfecdis.1c00387.

(37) Aslett, M.; Aurrecoechea, C.; Berriman, M.; Brestelli, J.; Brunk, B. P.; Carrington, M.; Depledge, D. P.; Fischer, S.; Gajria, B.; Gao, X.; Gardner, M. J.; Gingle, A.; Grant, G.; Harb, O. S.; Heiges, M.; Hertz-Fowler, C.; Houston, R.; Innamorato, F.; Iodice, J.; Kissinger, J. C.; Kraemer, E.; Li, W.; Logan, F. J.; Miller, J. A.; Mitra, S.; Myler, P. J.; Nayak, V.; Pennington, C.; Phan, I.; Pinney, D. F.; Ramasamy, G.; Rogers, M. B.; Roos, D. S.; Ross, C.; Sivam, D.; Smith, D. F.; Srinivasamoorthy, G.; Stoeckert, C. J. J.; Subramanian, S.; Thibodeau, R.; Tivey, A.; Treatman, C.; Velarde, G.; Wang, H. TriTrypDB: A Functional Genomic Resource for the Trypanosomatidae. Nucleic Acids Res. 2010, 38 (Database issue), D457–62. https://doi.org/10.1093/nar/gkp851.

(38) Spedale, G.; Timmers, H. T. M.; Pijnappel, W. W. M. P. ATAC-King the Complexity of SAGA during Evolution. Genes Dev. 2012, 26 (6), 527–541. https://doi.org/10.1101/gad.184705.111.

(39) Alonso, V. L.; Tavernelli, L. E.; Pezza, A.; Cribb, P.; Ritagliati, C.; Serra, E. Aim for the Readers! Bromodomains As New Targets Against Chagas’ Disease. Curr. Med. Chem. 2018. https://doi.org/10.2174/0929867325666181031132007.

(40) Tunyasuvunakool, K.; Adler, J.; Wu, Z.; Green, T.; Zielinski, M.; Žídek, A.; Bridgland, A.; Cowie, A.; Meyer, C.; Laydon, A.; Velankar, S.; Kleywegt, G. J.; Bateman, A.; Evans, R.; Pritzel, A.; Figurnov, M.; Ronneberger, O.; Bates, R.; Kohl, S. A. A.; Potapenko, A.; Ballard, A. J.; Romera-Paredes, B.; Nikolov, S.; Jain, R.; Clancy, E.; Reiman, D.; Petersen, S.; Senior, A. W.; Kavukcuoglu, K.; Birney, E.; Kohli, P.; Jumper, J.; Hassabis, D. Highly Accurate Protein Structure Prediction for the Human Proteome. Nature 2021, 596 (7873), 590–596. https://doi.org/10.1038/s41586-021-03828-1.

(41) Ritagliati, C.; Villanova, G. V.; Alonso, V. L.; Zuma, A. A.; Cribb, P.; Motta, M. C. M.; Serra, E. C. Glycosomal Bromodomain Factor 1 from Trypanosoma Cruzi Enhances Trypomastigote Cell Infection and Intracellular Amastigote Growth. Biochem. J. 2016, 473 (1), 73–85. https://doi.org/10.1042/BJ20150986.

(42) Alonso, V. L.; Villanova, G. V.; Ritagliati, C.; Motta, M. C. M.; Cribb, P.; Serra, E. C. Trypanosoma Cruzi Bromodomain Factor 3 Binds Acetylated α-Tubulin and Concentrates in the Flagellum during Metacyclogenesis. Eukaryot. Cell 2014. https://doi.org/10.1128/EC.00341-13.

(43) Villanova, G. V.; Nardelli, S. C.; Cribb, P.; Magdaleno, A.; Silber, A. M.; Motta, M. C. M.; Schenkman, S.; Serra, E. Trypanosoma Cruzi Bromodomain Factor 2 (BDF2) Binds to Acetylated Histones and Is Accumulated after UV Irradiation. Int. J. Parasitol. 2009. https://doi.org/10.1016/j.ijpara.2008.11.013.

(44) Rosón, J. N.; de Oliveira Vitarelli, M.; Costa-Silva, H. M.; Pereira, K. S.; da Silva Pires, D.; de Sousa Lopes, L.; Cordeiro, B.; Kraus, A. J.; Cruz, K. N. T.; Calderano, S. G.; Fragoso, S. P.; Siegel, T. N.; Elias, M. C.; da Cunha, J. P. C. Histone H2B.V Demarcates Strategic Regions in the Trypanosoma Cruzi Genome, Associates with a Bromodomain Factor and Affects Parasite Differentiation and Host Cell Invasion. bioRxiv 2021.

(45) Alsford, S.; Horn, D. Cell-Cycle-Regulated Control of VSG Expression Site Silencing by Histones and Histone Chaperones ASF1A and CAF-1b in Trypanosoma Brucei. Nucleic Acids Res. 2012. https://doi.org/10.1093/nar/gks813.

(46) Schulz, D.; Mugnier, M. R.; Paulsen, E. M.; Kim, H. S.; Chung, C. wa W.; Tough, D. F.; Rioja, I.; Prinjha, R. K.; Papavasiliou, F. N.; Debler, E. W. Bromodomain Proteins Contribute to Maintenance of Bloodstream Form Stage Identity in the African Trypanosome. PLoS Biol. 2015. https://doi.org/10.1371/journal.pbio.1002316.

(47) Lander, N.; Li, Z. H.; Niyogi, S.; Docampo, R. CRISPR/Cas9-Induced Disruption of Paraflagellar Rod Protein 1 and 2 Genes in Trypanosoma Cruzi Reveals Their Role in Flagellar Attachment. MBio 2015. https://doi.org/10.1128/mBio.01012-15.

(48) Taylor, M. C.; Kelly, J. M. PTcINDEX: A Stable Tetracycline-Regulated Expression Vector for Trypanosoma Cruzi. BMC Biotechnol. 2006. https://doi.org/10.1186/1472-6750-6-32.

(49) Alonso, V. L.; Ritagliati, C.; Cribb, P.; Serra, E. C. Construction of Three New Gateway® Expression Plasmids for Trypanosoma Cruzi. Mem. Inst. Oswaldo Cruz 2014. https://doi.org/10.1590/0074-0276140238.

(50) Nguyen Ba, A. N.; Pogoutse, A.; Provart, N.; Moses, A. M. NLStradamus: A Simple Hidden Markov Model for Nuclear Localization Signal Prediction. BMC Bioinformatics 2009. https://doi.org/10.1186/1471-2105-10-202.

(51) Goos, C.; Dejung, M.; Janzen, C. J.; Butter, F.; Kramer, S. The Nuclear Proteome of Trypanosoma Brucei. PLoS One 2017. https://doi.org/10.1371/journal.pone.0181884.

(52) Elias, M. C.; Marques-Porto, R.; Freymüller, E.; Schenkman, S. Transcription Rate Modulation through the Trypanosoma Cruzi Life Cycle Occurs in Parallel with Changes in Nuclear Organisation. Mol. Biochem. Parasitol. 2001, 112 (1), 79–90. https://doi.org/10.1016/s0166-6851(00)00349-2.

(53) Seal, J.; Lamotte, Y.; Donche, F.; Bouillot, A.; Mirguet, O.; Gellibert, F.; Nicodeme, E.; Krysa, G.; Kirilovsky, J.; Beinke, S.; McCleary, S.; Rioja, I.; Bamborough, P.; Chung, C.-W.; Gordon, L.; Lewis, T.; Walker, A. L.; Cutler, L.; Lugo, D.; Wilson, D. M.; Witherington, J.; Lee, K.; Prinjha, R. K. Identification of a Novel Series of BET Family Bromodomain Inhibitors: Binding Mode and Profile of I-BET151 (GSK1210151A). Bioorg. Med. Chem. Lett. 2012, 22 (8), 2968–2972. https://doi.org/10.1016/j.bmcl.2012.02.041.

(54) Picaud, S.; Wells, C.; Felletar, I.; Brotherton, D.; Martin, S.; Savitsky, P.; Diez-Dacal, B.; Philpott, M.; Bountra, C.; Lingard, H.; Fedorov, O.; Müller, S.; Brennan, P. E.; Knapp, S.; Filippakopoulos, P. RVX-208, an Inhibitor of BET Transcriptional Regulators with Selectivity for the Second Bromodomain. Proc. Natl. Acad. Sci. U. S. A. 2013, 110 (49), 19754–19759. https://doi.org/10.1073/pnas.1310658110.

(55) Picaud, S.; Leonards, K.; Lambert, J.-P.; Dovey, O.; Wells, C.; Fedorov, O.; Monteiro, O.; Fujisawa, T.; Wang, C.-Y.; Lingard, H.; Tallant, C.; Nikbin, N.; Guetzoyan, L.; Ingham, R.; Ley, S. V; Brennan, P.; Muller, S.; Samsonova, A.; Gingras, A.-C.; Schwaller, J.; Vassiliou, G.; Knapp, S.; Filippakopoulos, P. Promiscuous Targeting of Bromodomains by Bromosporine Identifies BET Proteins as Master Regulators of Primary Transcription Response in Leukemia. Sci. Adv. 2016, 2 (10), e1600760. https://doi.org/10.1126/sciadv.1600760.

(56) Meiler, J.; Baker, D. ROSETTALIGAND: Protein-Small Molecule Docking with Full Side-Chain Flexibility. Proteins Struct. Funct. Genet. 2006. https://doi.org/10.1002/prot.21086.

(57) Alonso, V. L.; Ritagliati, C.; Cribb, P.; Cricco, J. A.; Serra, E. C. Overexpression of Bromodomain Factor 3 in Trypanosoma Cruzi (Tc BDF3) Affects Parasite Differentiation and Protects It against Bromodomain Inhibitors. FEBS J. 2016, 283 (11), 2051–2066. https://doi.org/10.1111/febs.13719.

(58) Lander, N.; Chiurillo, M. A.; Docampo, R. Genome Editing by CRISPR/Cas9: A Game Change in the Genetic Manipulation of Protists. The Journal of eukaryotic microbiology. 2016. https://doi.org/10.1111/jeu.12338.

(59) Samworth, C. M.; Esposti, M. D.; Lenaz, G. Quenching of the Intrinsic Tryptophan Fluorescence of Mitochondrial Ubiquinol—Cytochrome-c Reductase by the Binding of Ubiquinone. Eur. J. Biochem. 1988. https://doi.org/10.1111/j.1432-1033.1988.tb13761.x.

