## Supplementary Figures for "Essential bromodomain *Tc*BDF2 as a drug target against Chagas disease"

^5^ Immunology Research Unit, Research, R&D GlaxoSmithKline, Gunnels Wood Road, Stevenage, Herts, SG1 2NY, UK.

^†^ Current address: Sciengement Lab Consulting, 28750 San Agustin del Guadalix, Spain.

**^#^** These Authors contributed equally to the manuscript.

**Supporting Information**

Supplementary Figure S1. Expression of *Tc*BDF2dm in epimastigotes.

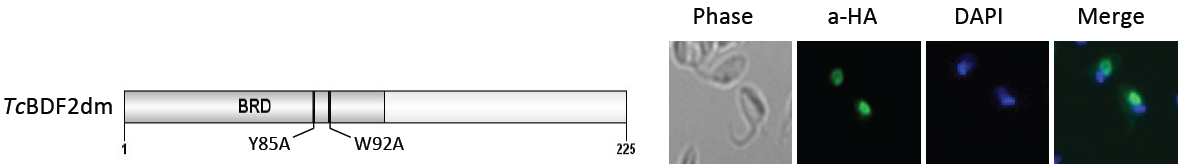

Supplementary Figure S2. Effect of over-expression of *Tc*BDF2 and *Tc*BDF2dm on the epimastigotes cell cycle measured by flow cytometry. Parasites overexpressing either *Tc*BDF2HA or *Tc*BDF2dm were subjected to flow cytometry to analyze the cell cycle progression at different times post tetracycline induction. Histograms are plotted as number of events vs. propidium iodide absorbance (PI-A).

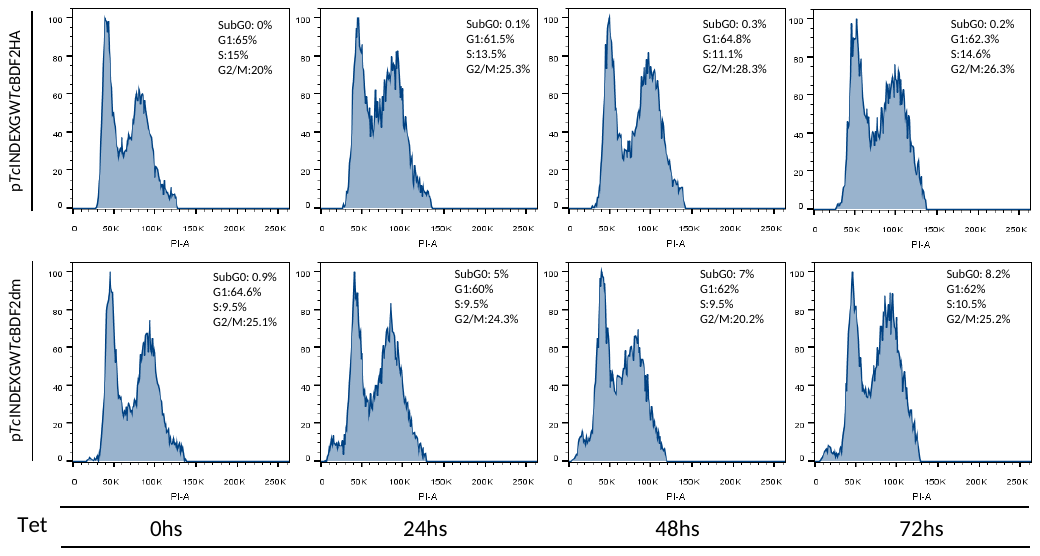

Supplementary Figure S3. Correct folding of recombinant *Tc*BDF2 and m*Tc*BDF2dm was corroborated by circular dichroism. Soluble proteins (5 mM) in 0.1 mM phosphate buffer, pH 8, were measured by circular dichroism spectroscopy using a spectropolarimeter Jasco J-810 (Easton, MD, USA).

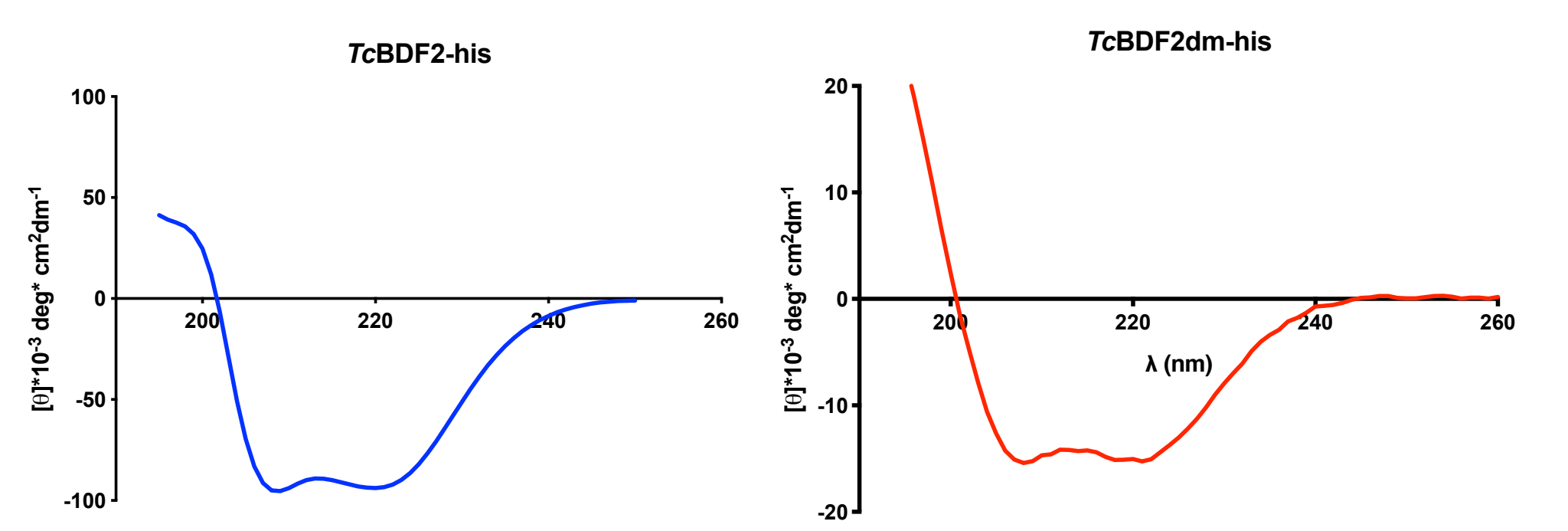

Supplementary Figure S4: Validation of RosettaLigand scripts and correlation with dissociation constants obtained by quenching of intrinsic tryptophan fluorescence. Predictions were made by extensive sampling of the conformational space using the RosettaLigand protocol. Briefly, the three-dimensional structures of the compounds were retrieved from PubChem database and their conformers were generated using BioChemicalLibrary suit (BCL). The structure of BD2 was generated with AlphaFold2 using the sequence of TcBDF2 from Dm28c strain. The starting pose of the compounds for local docking was obtained by aligning the co-crystal of BRD4 and JQ1(+) (PDB: 3MXF) with BD2 and taking the coordinates of the centroid of JQ1(+). Extensive sampling was carried out generating 7500 models for each compound, ordering them according to the interface energy calculated by Rosetta. Intermodel structural similarity was performed by calculating the ligand RMSD between each model and the lowest interface energy model. **A**: Comparison between the structure of BD2/BSP co-crystal (PDB: 6NIM) and the structures of the 5 lowest interface energy models. **B**: Funnel plot showing convergence of minimum energy models of BD2/BSP predictions in structures around 2.5 Å RMSD from the crystal structure (red circle). **C**: Box-plot comparing the energy and mean RMSD of the 5 lowest energy models for each prediction. BD2/BSP and BD2/iBET-151 displayed both the lowest median interface energy and the lowest Mean RMSD, consistent with the results obtained in vitro. **D**: Superposition of the 5 structures with the lowest interface energy for each compound. Obtaining minimum energy structures in heterogeneous poses is highly suggestive of a low affinity for the binding pocket, which is reflected in high RMSD values. **REU**: Rosetta Energy Units.

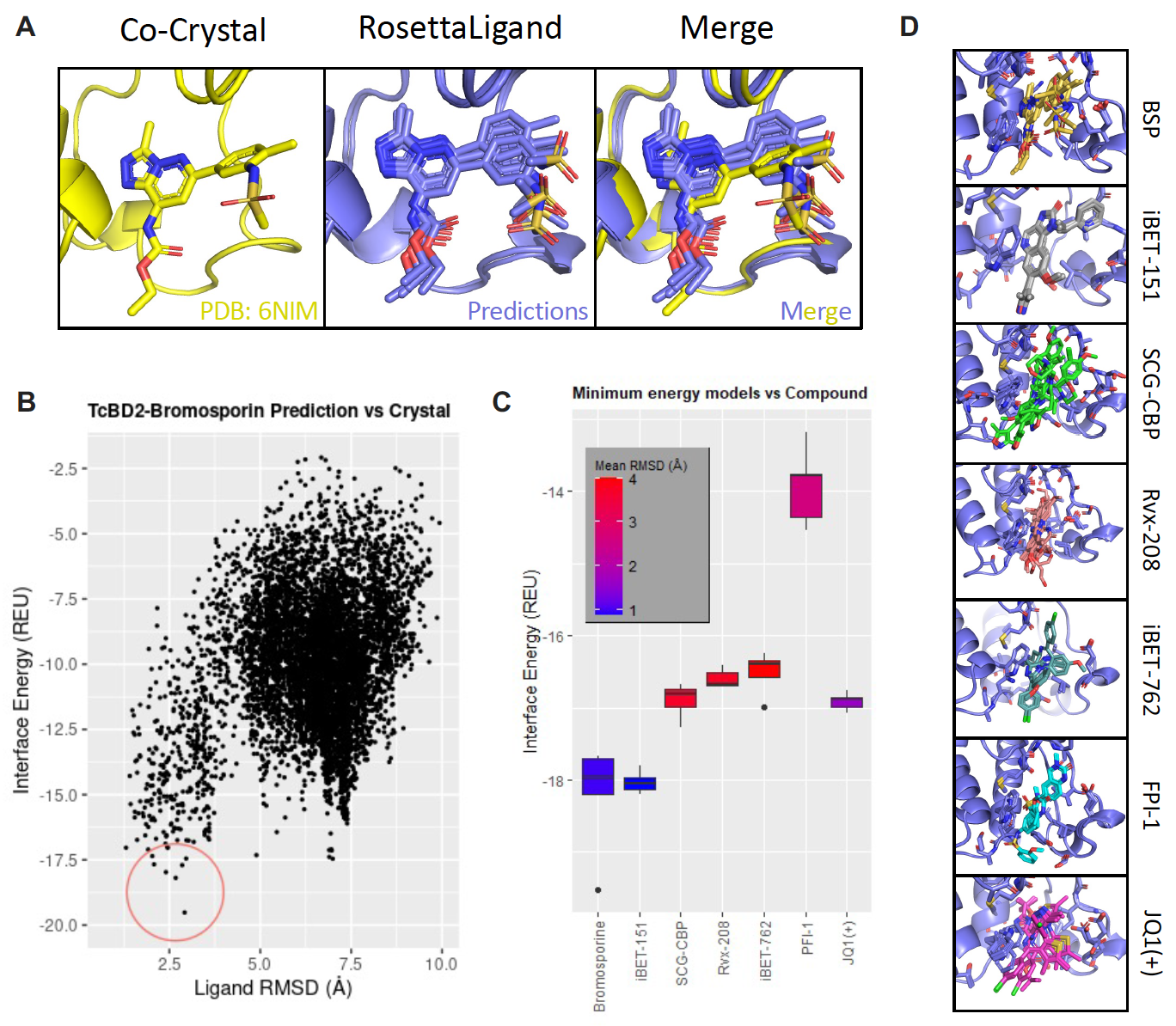
